# Not only perception but also grasping actions can obey Weber’s law

**DOI:** 10.1101/2022.06.15.496276

**Authors:** Zoltan Derzsi, Robert Volcic

## Abstract

Weber’s law, the principle that the uncertainty of perceptual estimates increases proportionally with object size, is regularly violated when considering the uncertainty of the grip aperture during grasping movements. The origins of this perception-action dissociation are debated and are attributed to various reasons, including different coding of visual size information for perception and action, biomechanical factors, the use of positional information to guide grasping, or, sensorimotor calibration. Here, we contrasted these accounts and compared perceptual and grasping uncertainties by asking people to indicate the visually perceived center of differently sized objects (Perception condition) or to grasp and lift the same objects with the requirement to achieve a balanced lift (Action condition). We found that the variability (uncertainty) of contact positions increased as a function of object size in both perception and action. The adherence of the Action condition to Weber’s law and the consequent absence of a perception-action dissociation contradict the predictions based on different coding of visual size information and sensorimotor calibration. These findings provide clear evidence that human perceptual and visuomotor systems rely on the same visual information and suggest that the previously reported violations of Weber’s law in grasping movements should be attributed to other factors.

## 1 Introduction

Weber’s law, one of the fundamental principles of perception, states that sensory precision decreases proportionally with the magnitude of a stimulus, or expressed differently, that the uncertainty of stimulus estimates increases with stimulus magnitude (Fechner, 1860). Examples of Weber’s law abound in all sensory modalities, with the exception of human grasping movements. When an object is about to be grasped, the variability in the maximum grip aperture between fingers is independent of object size, in contrast to the steady increase in variability with larger objects observed in perceptual estimates (Ganel et al., 2008). Although the finding that Weber’s law is violated in visually guided grasping actions is robust, less so is the agreement about its origin.

According to the Two-Visual Systems (TVS) hypothesis, perception and action rely on different computations of visual size information (Goodale, 2014; Goodale, 2011; Goodale & Milner, 1992). Vision-for-perception, mediated by the ventral visual stream, is based on relative metrics, taking into consideration not only the object size, but also general aspects of the visual scene. In contrast, vision-for-action, served by the dorsal visual stream, is based on absolute metrics reflecting the actual size of the object, which then leads to the violation of Weber’s law (Ganel et al., 2008, 2014). A second account, the biomechanical factors (BF) hypothesis, instead assumes that perception and action rely on the same visual size information and it attributes the dissociation to non-visual sources of noise that contaminate only the variability of the finger aperture in grasping (Bruno et al., 2016; Löwenkamp et al., 2015; Schenk et al., 2017; Uccelli et al., 2021; Utz et al., 2015). Weber’s law in grasping is violated, because, as object size increases, the finger opening is limited more and more by biomechanical factors, which artificially reduce the variability of the maximum grip aperture for larger objects. A third account, the digits-in-space (DIS) hypothesis, suggests that the maximum grip aperture is not influenced by visual size information at all, but it is instead an emergent property of the way the fingers are independently moved toward their contact positions on the object (Smeets & Brenner, 1999; Smeets et al., 2019; Verheij et al., 2012). If hand opening is influenced by positional information, and not size, it follows that the maximum grip aperture variability should not adhere to Weber’s law (Smeets & Brenner, 2008). A fourth account, the sensorimotor calibration (SMC) hypothesis, states that the dissociation between perception and action arises from differences in the availability of sensory feedback (Bingham & Mon-Williams, 2013; Bruno & Franz, 2009; Camponogara & Volcic, 2021; Cesanek et al., 2018; Schenk, 2012; Volcic & Domini, 2016, 2018). Visual and haptic feedback, which is missing during perceptual estimations, can only calibrate grasping actions and, by this, reduce the variability of the maximum grip aperture (Davarpanah Jazi et al., 2015; Heath et al., 2019).

To address this contentious debate, we explored another crucial component of successful grasping movements: the selection of optimal grasp contact positions on the surfaces of an object, which is essential to achieve a stable grasp (Goodale et al., 1994; Iberall et al., 1986; Klein et al., 2020; Kleinholdermann et al., 2013; Lederman & Wing, 2003). Stable grasps that minimize torques arising during lifting and prevent the object from slipping out of one’s grip are realized when the grasp axis passes as close as possible to the object’s center of mass. Specifically, we here introduce an experimental paradigm by which we pit the aforementioned hypotheses against each other by focusing on the *precision* with which grasp contact positions are selected on an object. In the Perception condition, we asked participants to indicate the visually perceived center of differently sized objects (Fig. 1A). In the Action condition, we asked participants to grasp and lift the same objects with the requirement to achieve a balanced lift (Fig. 1B). Thus, visual size information is essential in both conditions for an effective localization of the object center. The advantages of this paradigm are significant. First, by requiring the same grip aperture to grasp objects of all sizes we can avoid the potential constraints imposed by biomechanical factors. Second, we can evaluate grasping precision when the actual action goal is achieved and not ahead of it as it occurs when precision is measured at the time of the maximum grip aperture.

**Fig. 1.**
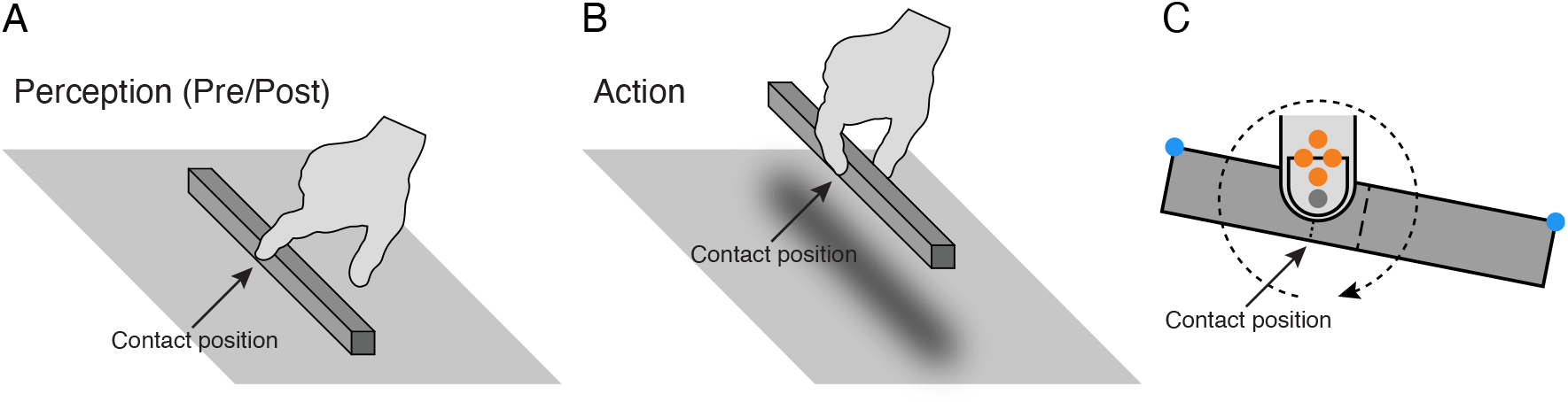
Methods. *A:* Perception condition: participants indicated the center of differently sized objects with their index finger, before (Pre) and after (Post) the Action condition. *B:* Action condition: participants grasped and lifted differently sized objects. The requirement was to achieve a balanced lift. *C:* Example of an off-center contact position. The position of the finger pad of the index finger was recorded with a virtual marker (gray point) with respect to four markers (orange points) attached on the distal phalanx of the index finger. Two markers (blue points) were attached on the edges of each object. The contact position (dotted line) was calculated with respect to the object center (dashed line). The circular dashed line represents the direction of object rotation and the tangential torque which provided visual and haptic feedback, respectively, whenever the object was lifted.

Determining the visually perceived center of an object can be seen as an alternative way of evaluating Weber’s law for object size. This is because the visually perceived center is based on the selection of a position on the object such that left and right segments of the object are not noticeably different from each other (Marshall & Halligan, 1989). If these judgments align with Weber’s law, the variability of the perceptual estimates of the object center should also exhibit a proportional increase with object size. This phenomenon has indeed been reported (Cattaneo et al., 2011; Manning et al., 1990; Nichelli et al., 1989; Riddoch & Humphreys, 1983; Wolfe, 1923). What is not yet known is whether and how does the variability of grasp contact positions depend on object size. If vision-for-action relies on absolute visual size information, as stated by the TVS hypothesis, we expect the grasping condition to violate Weber’s law also in this paradigm (Fig. 2A, solid line in mid panel). On the contrary, if Weber’s law is violated because of non-visual sources of noise (BF hypothesis), we expect the variability of the grasp contact positions to increase with object size with the same slope as in the perceptual condition, because biomechanical factors play no role here (Fig. 2A, long-dashed line in mid panel). Similarly, because the object center toward which fingers are steered can only be determined by considering the entire object size, the DIS hypothesis now predicts that the variability of grasp contact positions should adhere to Weber’s law (Fig. 2A, long-dashed line in mid panel). Lastly, if grasping movements are calibrated based on the visually perceived object rotation around the grasp axis and the haptically perceived tangential torques occurring whenever an object is grasped away from its center of mass (Fig. 1C), the SMC hypothesis predicts that the adherence to Weber’s law should be at least reduced, if not completely eliminated (Fig. 2A, dotted line in mid panel). In this case, the sensorimotor calibration might even affect perception, if the perceptual estimates are taken just after grasping (Fig. 2A, dotted line in right panel).

**Fig. 2.**
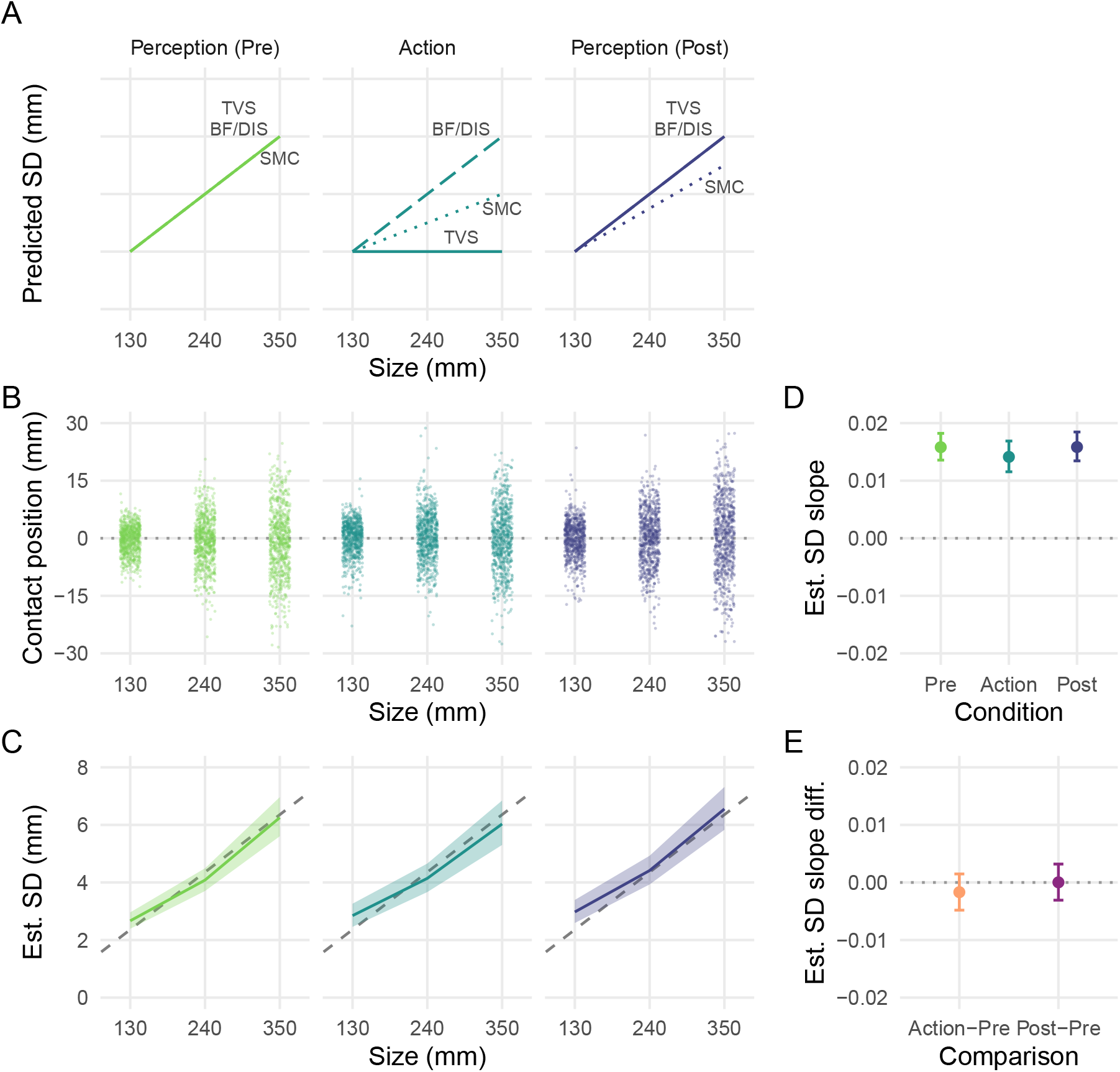
Predictions and results. *A:* Predicted SD as a function of size for the Two-visual systems (TVS), Biomechanical factors (BF), Digits-in-space (DIS), and Sensorimotor calibration (SMC) accounts in the Perception and Action conditions. *B:* Raw index finger contact positions relative to the center of each object. Points are slightly scattered horizontally for better visualization. *C:* Estimated SD as a function of size. Lines represent the mean and the shaded regions represent the 95% credible intervals of the posterior distributions of the Bayesian regression model. The dashed lines represent Weber’s law expressed as a straight line, with the slope (*k*) being the Weber fraction and the *y*-intercept being zero. A single Weber fraction (*k* ≈ .018) captures the increases in SD for all three conditions. *D:* Estimated slopes of the SD as a function of object size for each condition. Points represent the mean and the error bars the 95% credible intervals of the posterior distributions. *E:* Differences between estimated slopes of the SD for the Action*−*Pre and Post*−*Pre comparisons. Points represent the mean and the error bars represent the 95% credible intervals of the differences between posterior distributions.

## 2 Materials and Methods

### 2.1 Participants

Thirty-one naïve right-handed participants (age 19-35, 20 males) with normal or corrected-to-normal vision and no known history of neurological disorders participated in this study. They received subsistence allowance or course credit for their time. The study was undertaken with the understanding and informed written consent of each participant and the experimental procedures were approved by the Institutional Review Board of New York University Abu Dhabi.

### 2.2 Apparatus

Participants sat at a table with their right hand resting at a start position, aligned with their body midline. Stimuli objects were three 12 *×* 12 mm square-profile metal rods, 130 mm, 240 mm, and 350 mm in length, and they weighed 134 g, 256 g, and 382 g, respectively. The position of the index finger was recorded with sub-millimeter resolution by using an Optotrak Certus motion capture system (Northern Digital Inc., Waterloo, Ontario, Canada) controlled by the MOTOM toolbox (Derzsi & Volcic, 2018). Four infrared-light-emitting diodes (markers) were attached on the the distal phalanx of the index finger (Fig. 1C). The exact position of the index finger pad (virtual marker) was tracked by determining its location relative to the four markers. Two markers were attached on the edges of each object to track the position of its center (Fig. 1C). A pair of occlusion goggles (Red Scientific, Salt Lake City, UT, USA) was used to prevent vision between trials within each condition.

### 2.3 Procedure

The experiment consisted of three separate phases (72 trials each) performed in the following order: Perception (Pre) condition, Action condition, and Perception (Post) condition. In the Perception conditions, participants were asked to use their index finger to indicate the object center (Fig. 1A). In the Action condition, participants were asked to grasp the object with a thumb-index precision grip with vision of the hand and the object being constantly available (closed-loop grasping) and then lift the object while maintaining its horizontal balance (Fig. 1B). The Perception condition was performed twice: before (Pre) the Action condition to ensure that no visual or haptic feedback regarding the object true center location did influence perceptual estimates, and, after (Post) the Action condition to capture any effect of visual and haptic feedback on perceptual estimates.

Participants placed their hand at the start position before each trial, which could consist of indicating the object center or grasping and lifting the object depending on the condition. The experimenter then placed one of the three objects in front of the participant, 35 cm from the edge of the table and parallel to it. The position of the object was randomized in the range of *±*15 cm along this line, to avoid participants reaching always to the same location in space. The trial started when the goggles turned transparent. Once the participants indicated the object center (Perception conditions) or grasped and lifted the object (Action condition), the position of all the markers was recorded by the experimenter. Participants then placed the object down (in the Action condition) and returned the hand back to the start position. After this, vision was occluded in preparation for the next trial. Each of the three objects was presented 24 times in each condition (216 trials/participant), for a total of 6696 trials.

### 2.4 Data Analysis

All analyses were performed in R (R Core Team, 2022). The position of each contact position was obtained by performing an orthogonal projection of the fingertip virtual marker onto the line defined by the two markers positioned on the edges of each object. The contact position was then defined relative to the center of the object, with negative and positive values indicating contact positions to the left and to the right of the object center. Trials in which any of the markers were not captured correctly were discarded from further analysis (72 trials, approximately 1% in total).

Contact position data were analyzed using a Bayesian mixed-effects distributional regression model, estimated using the *brms* package (Bürkner, 2017; Carpenter et al., 2017). Unlike conventional regression models that assume a constant standard deviation across observations, a distributional regression model estimates not only the location parameters, but also the standard deviations of the location parameters. Our model included as predictors the categorical variable Condition (Perception (Pre), Action, Perception (Post)) and the continuous variable Size, which was centered before being entered in the model. This model estimated for each condition: the average contact position (contact position intercept), its change as a function of object size (contact position slope), the average standard deviation of the contact position (standard deviation of contact position intercept), and the standard deviation change of the contact position as a function of object size (standard deviation of contact position slope), the key measure in this study. The model included also independent random effects for participants to capture the dependency among data due to the repeated measures design. This statistical method allow us to simultaneously model all the raw data and directly estimate the main variable of interest (changes in standard deviation as a function of object size), in contrast to the multi-step approach normally used in which standard deviations are first calculated for each participant, size and condition separately, then fitted with linear models to extract the participants’ slopes, to finally pool them together to draw inferences.

Our model was fitted considering weakly informative prior distributions for each parameter to provide information about their plausible scale. We ran four Markov chains simultaneously, each for 4,000 iterations (1,000 warm-up samples) for a total of 12,000 post-warm-up samples. The obtained posterior distributions quantify the uncertainty about each estimated parameter conditional on the priors, model and data. We have summarized them by computing their means and their 95% credible intervals. To compare estimates between two conditions, we have computed the credible difference distributions by subtracting their respective posterior distributions, which were again summarized by computing their means and their 95% credible intervals.

### 2.5 Supplementary Material

Data and code are available at osf.io/e8c92/.

## 3 Results

Fig. 2B represents all the contact positions of the index finger relative to the center of each object in the Perception (Pre), Action and in the Perception (Post) conditions. These raw data clearly show that the contact positions were clustered around the center of each object, with no apparent systematic bias in either left (negative values) or right direction (positive values). The estimated average contact positions and their changes as a function of object size confirmed this observation; all posterior distributions of these estimates were centered on zero, equally extended in both positive and negative domains (contact position intercepts, Pre: *−*0.69 mm, 95% CI [*−*1.78, 0.41]; Action: 0.09 mm, 95% CI [*−*0.9, 1.1]; Post: 0.05 mm, 95% CI [*−*1.27, 1.36]) and did not depend on object size (contact position slopes, Pre: *−*0.003, 95% CI [*−*0.009, 0.004]; Action: *−*0.0004, 95% CI [*−*0.007, 0.006]; Post: 0.003, 95% CI

[*−*0.005, 0.010]).

The estimates of the standard deviation, the key measure used to compare the predictions made by the different hypotheses, are shown in Fig. 2C. The average standard deviations were of comparable magnitude in the three conditions (standard deviation of contact position intercepts, Pre: 4.08 mm, 95% CI [3.70, 4.49]; Action: 4.14 mm, 95% CI [3.67, 4.66]; Post: 4.41 mm, 95% CI [3.93, 4.93]).

Most importantly, it is evident that the standard deviations increased proportionally for larger object sizes (constant Weber fraction) not only in the Perception conditions, but also in the Action condition, contrary to the predictions of the TVS and SMC accounts (compare with Fig. 2A). In adherence to Weber’s law, the estimated slopes indicating the change in standard deviation of the contact positions as a function of object size (Pre: 0.016, Action: 0.014, Post: 0.016) and their 95% credible intervals were all positive and substantially above zero (Fig. 2D). No evidence was found of any difference in slopes between the Perception (Pre) and Action conditions, and, between the Perception (Post) and Perception (Pre) conditions. Both 95% credible intervals of the difference distributions overlapped with zero (Fig. 2E). Thus, both Perception and Action conditions conformed equally well to Weber’s law, showing no dissociation, but also no sign of sensorimotor calibration and transfer from the Action to the Perception (Post) condition. Computing the average slopes using the more common multi-step approach produces nearly identical slope values relating the standard deviations of contact positions and object size (Pre: 0.017, SD = 0.008; Action: 0.015, SD = 0.008; Post: 0.017, SD = 0.007).

## 4 Discussion and Conclusions

The main aim of this study was to investigate Weber’s law in relation to object size in both visual perception and visually guided grasping to address the debate about the origin of the perception-action dissociation. Specifically, we compared the uncertainty of perceptual estimates about the center of differently sized objects with the uncertainty with which optimal grasp contact positions are selected on those same objects, a crucial aspect of any visually guided grasping action (Goodale et al., 1994; Iberall et al., 1986; Klein et al., 2020; Kleinholdermann et al., 2013; Lederman & Wing, 2003). Our results show no sign of a perception-action dissociation and provide clear evidence that Weber’s law affects not only the variability of perceptual estimates, but also the variability of visually guided grasping as predicted by both the biomechanical factors (Bruno et al., 2016; Löwenkamp et al., 2015; Schenk et al., 2017; Uccelli et al., 2021; Utz et al., 2015) and the digits-in-space (Smeets & Brenner, 1999; Smeets et al., 2019; Verheij et al., 2012) hypotheses.

The finding that the variability of the contact positions in the Action condition increased with object size to the same degree as in the perceptual condition is not consistent with the TVS hypothesis (Ganel et al., 2008, 2014; Goodale, 2014; Goodale, 2011; Goodale & Milner, 1992). If vision-for-action were based on absolute metrics, we would have expected the variability in the Action condition to be constant over object sizes. This was clearly not the case, despite the fact that vision of the hand and the object was constantly available (closed-loop grasping), haptic feedback was provided by touching and lifting objects, and actions were directed toward real 3D objects. All these factors should have, if anything, favored the emergence of the perception-action dissociation. The range of sizes used for our objects was also not critical as similar ranges have been already used to study Weber’s law (Ganel et al., 2017; Hesse et al., 2021).

The present findings are also in contrast with the SMC hypothesis which predicted a reduction of the variability in the Action condition because of sensorimotor calibration, and, a potential transfer of this calibration on the perceptual estimates obtained just after the Action condition (Bingham & Mon-Williams, 2013; Bruno & Franz, 2009; Camponogara & Volcic, 2021; Cesanek et al., 2018; Schenk, 2012; Volcic & Domini, 2016, 2018). The precision of grasp contact positions definitely did not improve over successive repetitions even though inaccurate grasps provided direct visual and haptic feedback in the form of visually perceived object rotation and haptically perceived tangential torque which requires grip force adjustments to prevent rotational slip (Goodwin et al., 1998; Kinoshita et al., 1997). One possible reason for the lack of calibration could be linked to the slow adaptation that the grip force undergoes in the presence of torque load which might, in this particular case, negatively affect the ability to use haptic inputs effectively (Crevecoeur et al., 2011). Another reason that might have hindered or slowed down calibration was the randomized positioning of objects which required participants to assume slightly different arm postures for each grasping action.

Our study leads to an important implication about the role of Weber’s law in grasping actions: the fundamental assumption of the TVS hypothesis that perception and action rely on different computations of the visual input can only hold if we accept that absolute metrics is used exclusively for shaping the grip aperture during grasping, but the selection of optimal grasp contact positions is based on relative metrics. Thus, a recourse to a proposal involving two separate visual streams does not seem to be essential to explain how vision guides grasping movements. Instead, future studies will need to further explore how does the actual grasping precision occasionally conflate with factors intrinsic to motor behavior (e.g., Bhatia et al., 2022), how do task constraints determine whether grasping movements are guided by size or positional information, and, how does sensorimotor calibration affect grip apertures and optimal grasp contact positions over time.

## Funding

This work was partially supported by the NYUAD Center for Artificial Intelligence and Robotics, funded by Tamkeen under the NYUAD Research Institute Award CG010, and, by the NYUAD Center for Brain and Health, funded by Tamkeen under the NYUAD Research Institute Award CG012.

## Credit author statement

**Zoltan Derzsi:** Methodology, Software, Formal analysis, Validation, Writing - original draft; **Robert Volcic:** Conceptualization, Methodology, Formal analysis, Visualization, Validation, Data Curation, Writing - review and editing, Resources, Funding acquisition, Project administration, Supervision

## Notes

https://osf.io/e8c92/

### Competing Interest Statement

The authors have declared no competing interest.

### Summary of Updates

Introduction and Materials and Methods sections expanded with extra clarifications; Figure 2C revised.

## References

Bhatia, K., Löwenkamp, C., & Franz, V. H. (2022). Grasping follows Weber’s law: How to use response variability as a proxy for JND. Journal of Vision, 22 (12), 13. https://doi.org/10.1167/jov.22.12.13

Bingham, G. P., & Mon-Williams, M. A. (2013). The dynamics of sensorimotor calibration in reaching-to-grasp movements. Journal of Neurophysiology, 110 (12), 2857–2862. https://doi.org/10.1152/jn.00112.2013

Bruno, N., & Franz, V. H. (2009). When is grasping affected by the Müller-Lyer illusion? A quantitative review. Neuropsychologia, 47 (6), 1421–1433. https://doi.org/10.1016/j.neuropsychologia.2008.10.031

Bruno, N., Uccelli, S., Viviani, E., & De’Sperati, C. (2016). Both vision-for-perception and vision-for-action follow Weber’s law at small object sizes, but violate it at larger sizes. Neuropsychologia, 91, 327–334. https://doi.org/10.1016/j.neuropsychologia.2016.08.022

Bürkner, P.-C. (2017). brms: An R package for Bayesian Generalized Linear Mixed Models using Stan. Journal of Statistical Software, 80 (1), 1–28. https://doi.org/10.18637/jss.v080.i01

Camponogara, I., & Volcic, R. (2021). Integration of haptics and vision in human multisensory grasping. Cortex, 135, 173–185. https://doi.org/10.1016/j.cortex.2020.11.012

Carpenter, B., Gelman, A., Hoffman, M. D., Lee, D., Goodrich, B., Betancourt, M., Brubaker, M., Guo, J., Li, P., & Riddell, A. (2017). Stan: A probabilistic programming language. Journal of Statistical Software, 76 (1), 1–32. https://doi.org/10.18637/jss.v076.i01

Cattaneo, Z., Fantino, M., Tinti, C., Pascual-Leone, A., Silvanto, J., & Vecchi, T. (2011). Spatial biases in peripersonal space in sighted and blind individuals revealed by a haptic line bisection paradigm. Journal of Experimental Psychology: Human Perception and Performance, 37 (4), 1110–1121. https://doi.org/10.1037/a0023511

Cesanek, E., Campagnoli, C., Taylor, J. A., & Domini, F. (2018). Does visuomotor adaptation contribute to illusion-resistant grasping? Psychonomic Bulletin & Review, 25 (2), 827–845. https://doi.org/10.3758/s13423-017-1368-7

Crevecoeur, F., Giard, T., Thonnard, J.-L., & Lefevre, P. (2011). Adaptive control of grip force to compensate for static and dynamic torques during object manipulation. Journal of Neurophysiology, 106 (6), 2973–2981. https://doi.org/10.1152/jn.00367.2011

Davarpanah Jazi, S., Hosang, S., & Heath, M. (2015). Memory delay and haptic feedback influence the dissociation of tactile cues for perception and action. Neuropsychologia, 71, 91–100. https://doi.org/10.1016/j.neuropsychologia.2015.03.018

Derzsi, Z., & Volcic, R. (2018). MOTOM toolbox: MOtion Tracking via Optotrak and Matlab. Journal of Neuroscience Methods, 308, 129–134. https://doi.org/10.1016/j.jneumeth.2018.07.007

Fechner, G. T. (1860). Elemente der Psychophysik. Breitkopf und Härtel.

Ganel, T., Chajut, E., & Algom, D. (2008). Visual coding for action violates fundamental psychophysical principles. Current Biology, 18 (14), R599–601. https://doi.org/10.1016/j.cub.2008.04.052

Ganel, T., Freud, E., & Meiran, N. (2014). Action is immune to the effects of Weber’s law throughout the entire grasping trajectory. Journal of Vision, 14 (7). https://doi.org/10.1167/14.7.11

Ganel, T., Namdar, G., & Mirsky, A. (2017). Bimanual grasping does not adhere to Weber’s law. Scientific Reports, 7 (1). https://doi.org/10.1038/s41598-017-06799-4

Goodale, M. A. (2014). How and why the visual control of action differs from visual perception. Proceedings of the Royal Society B: Biological Sciences, 281 (1785), 20140337–20140337. https://doi.org/10.1098/rspb.2014.0337

Goodale, M. A. (2011). Transforming vision into action. Vision Research, 51 (13), 1567–1587. https://doi.org/10.1016/j.visres.2010.07.027

Goodale, M. A., Meenan, J. P., Bülthoff, H. H., Nicolle, D. A., Murphy, K. J., & Racicot, C. I. (1994). Separate neural pathways for the visual analysis of object shape in perception and prehension. Current Biology, 4 (7), 604–610. https://doi.org/10.1016/S0960-9822(00)00132-9

Goodale, M. A., & Milner, A. D. (1992). Separate visual pathways for perception and action. Trends in Neurosciences, 15 (1), 20–25. https://doi.org/10.1016/0166-2236(92)90344-8

Goodwin, A. W., Jenmalm, P., & Johansson, R. S. (1998). Control of grip force when tilting objects: Effect of curvature of grasped surfaces and applied tangential torque. Journal of Neuroscience, 18 (24), 10724–10734. https://doi.org/10.1523/JNEUROSCI.18-24-10724.1998

Heath, M., Chan, J., & Davarpanah Jazi, S. (2019). Tactile-based pantomime grasping: Knowledge of results is not enough to support an absolute calibration. Journal of Motor Behavior, 51 (1), 10–18. https://doi.org/10.1080/00222895.2017.1408559

Hesse, C., Harrison, R. E., Giesel, M., & Schenk, T. (2021). Bimanual grasping adheres to Weber’s law. i-Perception, 12 (6), 204166952110545. https://doi.org/10.1177/20416695211054534

Iberall, T., Bingham, G. P., & Arbib, M. A. (1986). Opposition space as a structuring concept for the analysis of skilled hand movements. In H. Heuer & C. Fromm (Eds.), Generation and modulation of action patterns (pp. 158–173). Springer-Verlag.

Kinoshita, H., Bäckström, L., Flanagan, J. R., & Johansson, R. S. (1997). Tangential torque effects on the control of grip forces when holding objects with a precision grip. Journal of Neurophysiology, 78 (3), 1619–1630. https://doi.org/10.1152/jn.1997.78.3.1619

Klein, L. K., Maiello, G., Paulun, V. C., & Fleming, R. W. (2020). Predicting precision grip grasp locations on three-dimensional objects. PLOS Computational Biology, 16 (8), e1008081. https://doi.org/10.1371/journal.pcbi.1008081

Kleinholdermann, U., Franz, V. H., & Gegenfurtner, K. R. (2013). Human grasp point selection. Journal of Vision, 13 (8). https://doi.org/10.1167/13.8.23

Lederman, S., & Wing, A. (2003). Perceptual judgement, grasp point selection and object symmetry. Experimental Brain Research, 152 (2), 156–165. https://doi.org/10.1007/s00221-003-1522-5

Löwenkamp, C., Gärtner, W., Haus, I. D., & Franz, V. H. (2015). Semantic grasping escapes Weber’s law. Neuropsychologia, 70, 235–245. https://doi.org/10.1016/j.neuropsychologia.2015.02.037

Manning, L., Halligan, P. W., & Marshall, J. C. (1990). Individual variation in line bisection: A study of normal subjects with application to the interpretation of visual neglect. Neuropsychologia, 28 (7), 647–655. https://doi.org/10.1016/0028-3932(90)90119-9

Marshall, J. C., & Halligan, P. W. (1989). When right goes left: An investigation of line bisection in a case of visual neglect. Cortex, 25 (3), 503–515. https://doi.org/10.1016/S0010-9452(89)80065-6

Nichelli, P., Rinaldi, M., & Cubelli, R. (1989). Selective spatial attention and length representation in normal subjects and in patients with unilateral spatial neglect. Brain and Cognition, 9 (1), 57–70. https://doi.org/10.1016/0278-2626(89)90044-4

R Core Team. (2022). R: A language and environment for statistical computing. R Foundation for Statistical Computing. https://www.R-project.org/

Riddoch, M., & Humphreys, G. W. (1983). The effect of cueing on unilateral neglect. Neuropsychologia, 21 (6), 589–599. https://doi.org/10.1016/0028-3932(83)90056-8

Schenk, T. (2012). No dissociation between perception and action in patient DF when haptic feedback is withdrawn. Journal of Neuroscience, 32 (6), 2013–2017. https://doi.org/10.1523/JNEUROSCI.3413-11.2012

Schenk, T., Utz, K. S., & Hesse, C. (2017). Violations of Weber’s law tell us more about methodological challenges in sensorimotor research than about the neural correlates of visual behaviour. Vision Research, 140, 140–143. https://doi.org/10.1016/j.visres.2017.05.017

Smeets, J. B. J., & Brenner, E. (1999). A new view on grasping. Motor Control, 3 (3), 237–271. https://doi.org/10.1123/mcj.3.3.237

Smeets, J. B. J., & Brenner, E. (2008). Grasping Weber’s law. Current Biology, 18 (23), R1089–R1090. https://doi.org/10.1016/j.cub.2008.10.008

Smeets, J. B. J., van der Kooij, K., & Brenner, E. (2019). A review of grasping as the movements of digits in space. Journal of Neurophysiology, 122 (4), 1578–1597. https://doi.org/10.1152/jn.00123.2019

Uccelli, S., Pisu, V., & Bruno, N. (2021). Precision in grasping: Consistent with Weber’s law, but constrained by “safety margins”. Neuropsychologia, 163, 108088. https://doi.org/10.1016/j.neuropsychologia.2021.108088

Utz, K. S., Hesse, C., Aschenneller, N., & Schenk, T. (2015). Biomechanical factors may explain why grasping violates Weber’s law. Vision Research, 111, 22–30. https://doi.org/10.1016/j.visres.2015.03.021

Verheij, R., Brenner, E., & Smeets, J. B. J. (2012). Grasping kinematics from the perspective of the individual digits: A modelling study. PLoS ONE, 7 (3), e33150–e33150. https://doi.org/10.1371/journal.pone.0033150

Volcic, R., & Domini, F. (2016). On-line visual control of grasping movements. Experimental Brain Research, 234 (8), 2165–2177. https://doi.org/10.1007/s00221-016-4620-x

Volcic, R., & Domini, F. (2018). The endless visuomotor calibration of reach-to-grasp actions. Scientific Reports, 8 (1), 14803. https://doi.org/10.1038/s41598-018-33009-6

Wolfe, H. K. (1923). On the estimation of the middle of lines. The American Journal of Psychology, 34 (3), 313–358. https://doi.org/10.2307/1413954

